# Structural Investigation of Oat Protein Isolate in Aqueous Medium by using Synchrotron Small-angle X-ray Scattering

**DOI:** 10.1101/2020.11.14.363911

**Authors:** Ji Li, Qingrong Huang

**Affiliations:** Nutraceutical Corporation, 1777 Sun Peak Dr., Park City, Utah 84098; Department of Food Science, Rutgers University, 65 Dudley Road, New Brunswick, New Jersey 08901-8520, U.S.A

**Keywords:** Oat protein isolate (OPI), synchrotron small-angle X-ray scattering, form factor, effective structure factor, *ab initio* reconstitution

## Abstract

Oat protein isolate (OPI) is among the plant proteins with valuable functionalities (e.g. emulsification) during daily supplement intake. Understanding their structures helps to manipulate oat proteins at small scale, which enables the appropriate deployment of their functions. Based upon such understanding, the molecular structure of oat protein isolate (OPI) in aqueous medium was investigated by synchrotron small-angle X-ray scattering (SAXS), and this study allows a structural reconstitution of OPI in aqueous medium. Besides, this SAXS study is complimentary to the previous study (Liu et al. *J. Agric. Food Chem*. 2009, 57, 4552–4558)^1^. From form factor fitting, we confirmed that OPI aqueous solutions at low concentrations (0.3~2 mg/mL) obtained a disk conformation (41.4×41.4×10.2 Å^3^). Once protein concentration increased to 5 mg/mL and 10 mg/mL, the individual disk proteins formed large-dimensional rodlike aggregates, which was evidenced by the analyses of effective structure factor and pair distribution function (PDF). Based on the PDF results, the *ab initio* models of OPI particles at low concentrations (0.3 mg/mL to 2.0 mg/mL) were restored by using GASBOR algorithm. Finally, we found that weak attraction between OPI particles occurred, which was verified by second virial coefficient and pair potential.

## 1. Introduction

Cereal proteins are helpful for maintaining regular health of human body. In many developing countries, the consumption of cereal products even occupies 70% to 90% of the total protein consumption. Typical cereal proteins are prolamines such as gluten^2^ and zein^3^. Under the family of cereal proteins, oat proteins are the only species which contains a globulin or legume-like protein, avenalin, as the major (80%) storage protein^4^. A small portion of oat proteins is avenin, a prolamine. Oat proteins are known for high nutritional quality which can be equivalent to proteins from soy, meat, milk, and egg^5,6^. With its amphiphilic nature, oat protein is endowed with interfacial functionalities such as emulsification7, water holding8, and foaming9. Recently, Xu et al. found that oat protein could ameliorate exercise-induced fatigue by observing the levels of liver glycogen, blood urea nitrogen, malondialdehyde in serum, and activities of lactic dehydrogenase and superoxide dismutase in Kun-ming mice model^10^.

The proteins’ functionalities are directly-related to their structures at micro/nano-levels. Unveiling protein structures in solution enables us to understand protein folding^11^, interaction between protein and active compounds such as polyphenols^12,13^, and predictable protein conformation or behavior^14,15^. Various approaches have been adopted to restore the protein structures at different levels. Proteins’ secondary structures such as α-helix, β-sheet, and random coil are notably predicted locally via ultraviolet-circular dichroism (UV-CD)^16^ and infrared spectroscopy (IR)^17^. Both approaches have an upper limit on the accuracy of secondary structure prediction owing to their failure to account for nonlocal interaction. The prediction of local structure partially discloses the whole protein structure, and still lack of higher order structural information such as tertiary, quaternary structures. With the knowledge of protein sequences, proteins’ spatial structures can be estimated by comparing the sequence identity to known proteins^18^ or searching the protein conformation with the lowest free energy^19^. Although those two computational approaches have been widely utilized to realize protein restoration^20^, those approaches have limitations of amino acid sequence identity and computational resource.

Small-angle X-ray scattering (SAXS) with advanced synchrotron irradiation is a powerful tool for probing plant proteins’ structures in different mediums^21^. The synchrotron facilities bear the advantages of quick measurement, sample protection, and complementation to other characterizations. With the flexibility of measurement conditions, SAXS becomes a useful alternative, allowing the investigation of proteins in different conditions. Besides, *ab initio* modeling based on pair distribution function (PDF) analysis of SAXS data can quickly perform the restoration of low-resolution protein envelopes. Several algorithms are available to restore low-resolution protein shape directly from the scattering profiles, including CRYSOL^22^, DAMMIN^23^, and GASBOR^24^. In detail, *ab initio* protein shapes can be restored by using a dummy residue representation of the protein model which describes the protein internal structure more adequately^25^. Different from beads in DAMMIN, dummy residues in GASBOR correspond to amino acid residues which are closer to the scattering of protein in solution. Meanwhile, the positions of those dummy residues are still determined by the simulated annealing-driven minimization to fit the experimental SAXS. In the case of crop protein Z19 α-zein, Forato et al. performed GASBOR algorithm for SAXS profile of Z19 α-zein in 90% ethanol/water solution^26^, and the restored low-resolution model confirmed its extended structure with 12 by 130 Å, which was in agreement with the model proposed by Matsushima et al^27^.

In this article, we continue the structural investigation of oat protein isolate (OPI) in aqueous medium based on our previous study^1^. In the past, we quantified the secondary structure of OPI with the attentuated total reflectance-Fourier transform infrared spectroscopy (ATR-FTIR), and observed the disk-like conformation of OPI through tapping mode-atomic force microscopy (TP-AFM). Synchrotron small-angle X-ray scattering was used to analyze the protein conformation in aqueous medium, which is complementary to the previous structural study in solid state. The solubilities of OPI at different pHs give the working window for SAXS study. Form factors of solid sphere, rod, and ellipsoid are used to fit the SAXS profiles of OPI in aqueous medium. The dimension of individual OPI particle is determined by form factor fitting and Guinier analysis, while the aggregation behavior is semi-quantified by the effective structure analysis. Pair distribution function (PDF) is employed to give the size distribution of OPI in aqueous medium. Based on PDF, *ab initio* restoration of OPI in aqueous medium is obtained by using the GASBOR algorithm

## 2. Materials and methods

### 2.1. Materials

Oat protein isolate (OPI) was isolated by isoelectric precipitation method according to our previous work^1^. Initially, 100 g oat flour was mixed with 600 g water, while the solution pH was adjusted to 10.0 by using 2 M NaOH. After fully dissolution, the blended slurry was filtered through wiremeshes of 100 μm, 75 μm, and 50 μm. Those filtered fine slurries were centrifuged at 3000 g for 15 min at ambient environment. The first-round supernatant was precipitated at oat proteins’ isoelectric points after pH was adjusted to 5.0 by using 0.5 M HCl. Then, the oat protein samples were subject to centrifugation at 3000g for another 15min. The obtained oat protein pellet was washed with DI water for 3 times. Finally, the OPI powder was lyophilized by an LG-5 freeze dryer (Shanghai Centrifuge Institute Co., Ltd., China). Milli-Q water (18.3 Ω) was used in all experiments.

### 2.2. Protein solubility

The determination of protein solubility was modified based on the method of Wang et al^28^. Oat protein isolate samples were dispersed in Milli-Q water at 1% (w/v), and the pH value was adjusted between 2.0 and 10.0 with 0.5 M HCl and 0.5 M NaOH. After 30 min stirring, pH-adjusted protein sample was centrifuged at 10,000 g for 10 min at 20 °C. The protein concentrations of supernatant were measured by Lowry’s method with bovine serum albumin (BSA) as the standard. The protein solubility was described as grams of soluble protein/ 100 g of protein sample.

### 2.3. Synchrotron small-angle X-ray scattering (SAXS)

SAXS profiles were collected by using the 18ID undulator beamline of the Biophysics Collaborative Access Team at the Advanced Photon Source, Argonne National Laboratory^29^. The sample-to-detector distance was set at 2.3 meter to cover a Q range of 0.006-0.37 Å. The experimental set-up also includes a 3 m sample-to-detector-length camera and another 0.3 m sample-to-detector-length camera with the high-sensitivity CCD detector. A flow cell of 1.5 mm diameter capillary equipped with a brass block (thermostatted with a water bath) was utilized to hold samples. In order to minimize the radiation damage, a MICROLAB 500 Hamilton pump was applied to load samples to the flow cell at a constant rate (10 mL/s). The X-ray wavelength was 1.033 Å and a short exposure period of 1 s was used to acquire the scattering data. The whole experiment was kept at room temperature. Fifteen curves were collected for each sample and their averaged curves were utilized for further analysis. The final SAXS profiles were gained after subtracting the solvent background. The angular scale was calibrated using the scattering peaks of silver behenate.

### 2.4. SAXS analysis

Synchrotron small-angle X-ray scattering is used here to study the structure and interaction of biopolymers in solution^30^. The total scattering intensity, I(Q), for a monodisperse protein can be expressed as^31,32^:

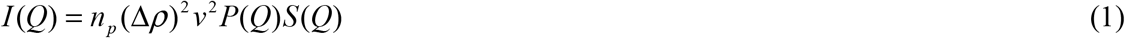

where *n*_p_ is the number density of colloidal particles per unit volume in the solution, *v* is the specific volume of protein which can be predicted through the empirical equation proposed by Fischer et al.,^33^ *Q=(4πX)sin(θ/2)* is the scattering vector, where λ is the wavelength of the X-ray beam, *θ* is the scattering angle, Δρ is the contrast of electron densities between proteins and solvent, *P(Q)* is the form factor of a given protein which reflects the shape of individual protein, and *S(Q)* is the structure factor which suggests the protein aggregation.

For a solid sphere particle with a radius *R,* the form factor can be described as^32^:

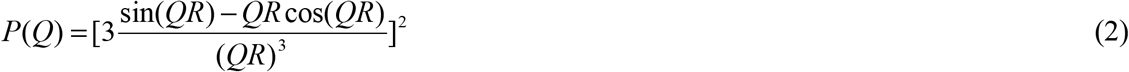

For an ellipsoidal particle with a parallel radii of the equivalent ellipsoid of the protein, *a* and a perpendicular radii of the equivalent ellipsoid of the protein, *b*, the form factor can be expressed^34,35^ as:

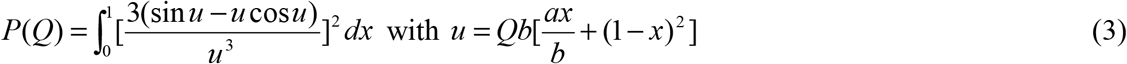

For a circular disk with disk radius *R* and disk thickness *L*, the form factor is shown as^36^:

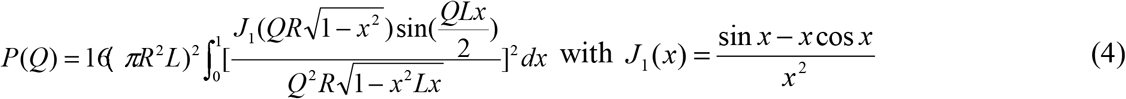

The protein aggregation and protein-protein interaction upon concentration increase in solutions can be described by using the effective structure factor^37^ *S*(Q,*C*) which is written as^38^:

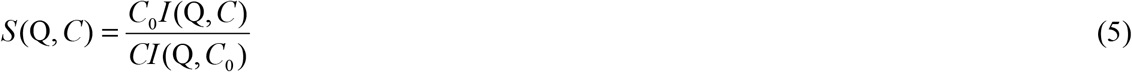

Here I(Q,*C*) is the scattering intensity profile from a protein solution with concentration *C*, and *C*_0_ corresponds to dilute solution in which protein aggregation is negligible.

Other analytical tools such as Guinier analysis and pair distribution function (PDF) were used to obtain the radii of gyration *R_g_,* forward scattering I(0), and size distribution. The Guinier approximation is given by:

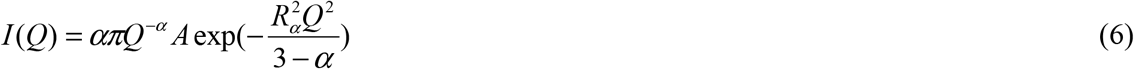

where α=0 for solid sphere; α=1 for rod-like object; and α=2 for sheet-like object. When classic Guinier fit (α=0) is performed, *A* is equal to I(0), the scattering intensity at Q=0, and R_g_, the radius of gyration, is equal to R_α_. When rod-like Guinier fit is performed, R_c_, the cross-section radius of gyration, is equivalent to 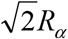. When sheet-like Guinier fit is performed, T, the thickness, is equivalent to 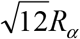. During fitting, the first and last few points from the fitted Guinier curves were changed by appropriate handlers. GNOM package was used to generate the pair distribution function (PDF) *P(r)* which displayed the probability of finding one point within the particle dimension D_m_ at a distance *r* from a given point^39^. PDF shows size distribution and shape of particle envelope^40^.

### 2.5. *ab initio* restoration

The low-resolution structures (2-3 nm^-1^) of oat protein isolate (OPI) in aqueous medium were restored by using algorithm GASBOR. GASBOR generates the configuration as an assembly of dummy residues. The simulated annealing is also employed to provide a chain compatible spatial distribution of dummy residues inside the same search volume of a diameter D_max_. The simulation input is the GNOM file or the PDF analyzed file. 200 to 500 number of dummy residues, reciprocal space mode, and real space mode were all used during simulation.

## 3. Results and discussion

### 3.1. Protein solubility

Protein solubility (PS) in aqueous medium is the prerequisite for numerous applications of oat protein isolate (OPI). **Figure 1** exhibits the PS profile of OPI within the pH range from 2.0 to 10.0. The PS reached minimum at pH 5.0, and gradually increased when pH increased from 5.0 to 10.0 or decreased from 5.0 to 2.0. Similar U-shape PS profile can also be found in other crop proteins such as globulin from Mungbean ^41^ and rice bran protein^42^. The PS profile is related to the molecular interactions between proteins, especially electrostatic interaction. At pH 5.0, OPI is in neutral status, and the protein surface charges have been screened. Under such circumstance, oat protein easily aggregated together to precipitate out. At pH 2.0-5.0 and pH 5.0-10.0, oat protein carries positive charges and negative charges, respectively. Besides, the surface charge density is pH-dependent as well. The electrostatic interaction (i.e. electrostatic repulsion) between oat proteins ensures the well-dispersed proteins in the medium, and the stronger electrostatic repulsion, the better dispersibility of protein particles in medium. With the knowledge of PS profile, the working window for the structure investigation of OPI in medium is determined to be within pH 6.5-7.0, which meets the physiological condition of human body.

**Figure 1.**
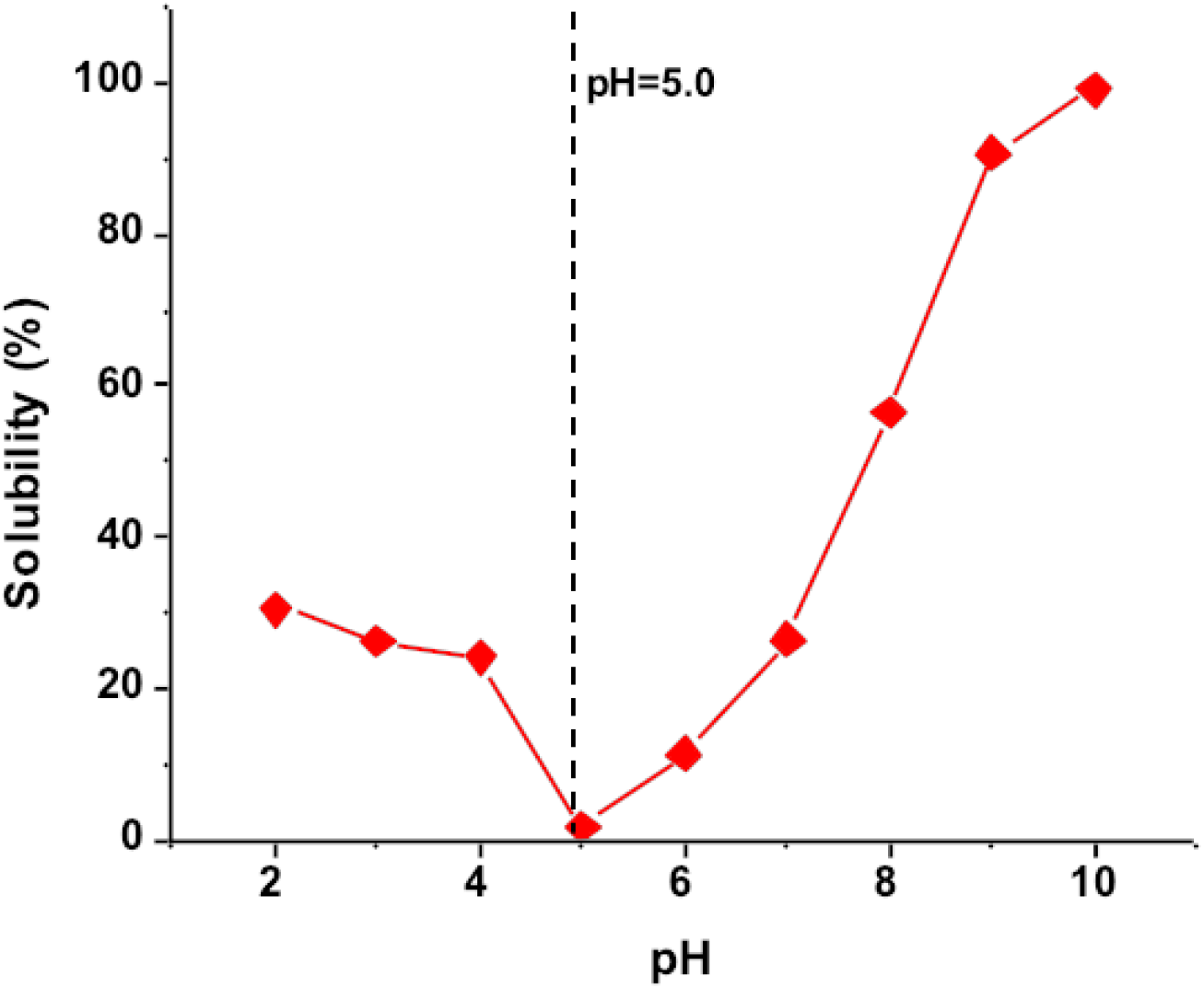
Solubility profile of oat protein isolate (OPI) extracted by isoelectric precipitation method

### 3.2. Protein conformation

In a dilute solution, the total scattering intensity is the sum scattering of total individual protein particles. The protein aggregation can be negligible under that condition, which facilitates the study of the shape of individual protein particle. The small-angle X-ray (SAXS) profile of 1 mg/mL oat protein isolate (OPI) was fitted by different form factors, including solid sphere, ellipsoidal, and circular disk, which is shown in **Figure 2A**. The fitted data from three form factors matched the experimental scattering intensity well for Q < 0.06 Å^−1^, meanwhile, the solid sphere form factor fitted the curve best only up to 0.078 Å^-1^ and the fitting of oblate ellipsoid form factor covered a more lengthy Q range (up to 0.13 Å^-1^). The circular disk form factor fitted the curve best within the whole scattering range. The dimensions from form factor fittings of OPI at 1 mg/mL are displayed in **Table 1**. The averaged radius *r* and thickness *L* of disk-like protein particle are 41.4 Å and 10.2 Å, respectively (**Table 1**). Based on the trials of solid sphere, ellipsoidal, and circular disk form factor fittings, we applied the circular disk form factor to the SAXS profiles of OPI solutions within a concentration range of 0.3-10 mg/mL (**Figure 2B**). All fittings of circular disk form factor performed well in the whole Q range. Thus, it again suggested that circular disk gIve the best description for the conformation of OPI in aqueous medium. The fitted dimensions, cross-section radius of disk (*r*) and disk thickness (*L*), are shown in **Figure 2C**. Those two parameters show little concentration-dependence, indicating the stability of circular disk form factor fitting.

**Figure 2.**
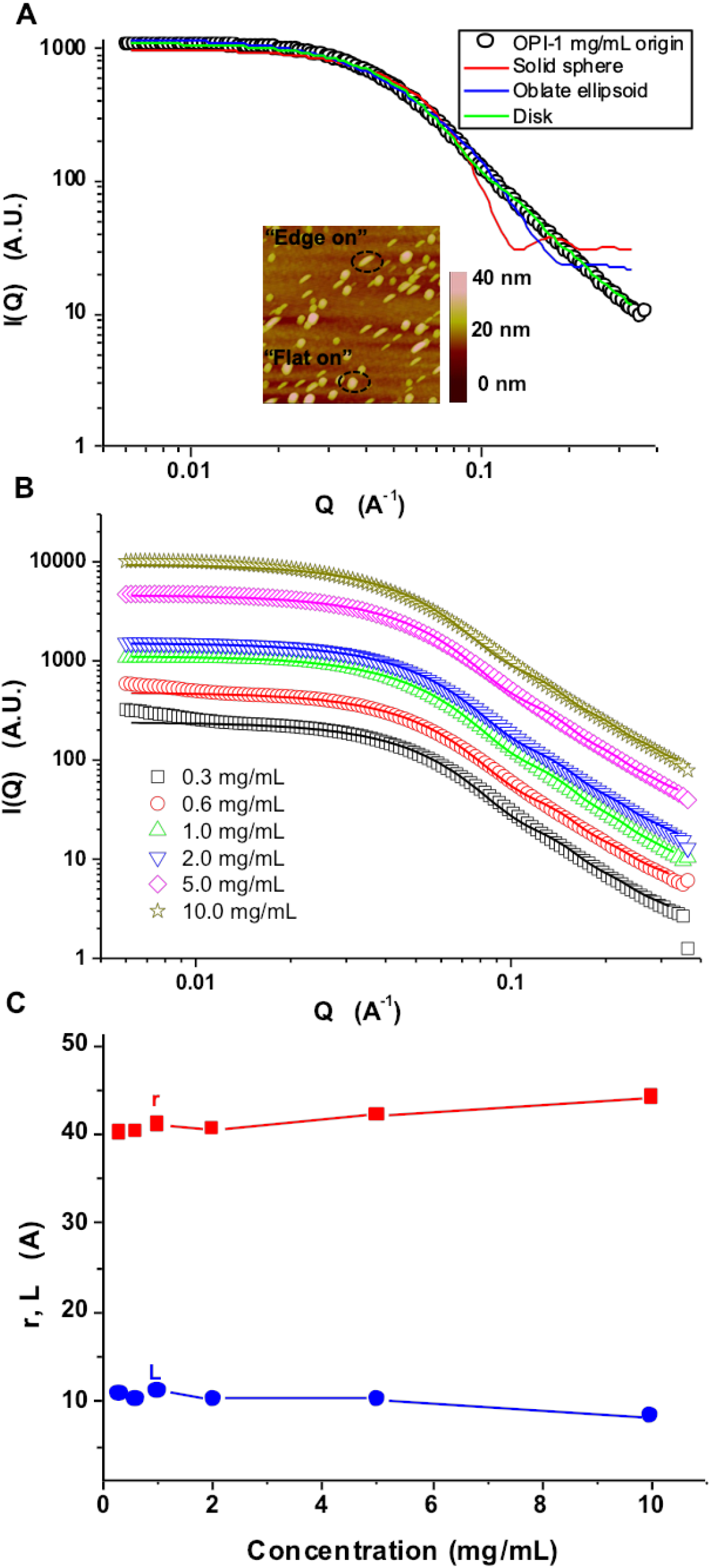
(A) Small-angle X-ray scattering (SAXS) profile and form factor fittings for 1 mg/mL oat protein isolate (OPI) aqueous solution; (B) SAXS profile and circular disk form factor fittings of 0.3-10 mg/mL OPI aqueous solutions; (C) Radius of disk, *r* and thickness of disk, *L* obtained from circular disk form factor fitting. The previous tapping mode-atomic force microscopy 2 μm × 2 μm height image of 0.5 mg/mL OPI solution on silicon wafer was also presented.

**Table 1.** Form factor fitting for oat protein isolate (OPI) at 1 mg/mL

| Form factor | Dimension |
| --- | --- |
| Solid sphere | $r = 33 \pm 0.0 \text{ \AA}$ |
| Oblate ellipsoid | $a = 20 \pm 0.0 \text{ \AA}, b = 72 \pm 0.0 \text{ \AA}$ |
| Circular disk | $r = 41.4 \pm 0.0 \text{ \AA}, L = 10.2 \pm 0.1 \text{ \AA}$ |

Previously, we observed the “edge on” disk and the “flat on” disk by using tapping mode-atomic force microscopy (AFM) (**Figure 2A inset**)^43^. The “edge on” disk meant that the disk edge was toward our eye perspective, while “flat on” disk suggested that the circular surface of the disk is toward our eye perspective. The circular disk form factor fitting of OPI is in agreement with our previous AFM results.

### 3.3. Guinier analysis

Guinier approximations for solid sphere, rod-like and sheet-like object were all used to treat the SAXS profiles of 0.3-10 mg/mL OPI aqueous solutions. The fitting performance of solid sphere and sheet-like Guinier analysis is shown in **Figure 3A** and **Figure 3B**. The appropriate Q range was selected to ensure the Q*R_g_<1.3. The radius of gyration R_g_ and thickness T were extracted from the Guinier plot, and the concentration dependence of R_g_, T, and 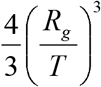 was presented in **Figure 3C**. The T was a concentration-independent, and was 10.5 Å at all concentrations, which aligned with the thickness from disk form factor fitting (**Figure 2C**). Simultaneously, R_g_ kept at 30 Å within the concentration range of 0.3-2 mg/mL, while increased to 34 Å once concentration reached 10 mg/mL. The axial ratio determined through 4/3(R_g_/T)^3^ was approximately 30 at low concentration (0.3-2.0 mg/mL). Unlike α-zein^34^, oat protein displayed an increase of axial ratio upon higher concentration, and the axial ratio increased to 37 at 5.0 mg/mL and 47 at 10.0 mg/mL, which suggested an protein growth in a uniaxial direction. The pronounced increase of R_g_ and axial ratio was caused by the formation of agglomerate through protein stacking. The rodlike Guinier analysis was performed as well. The cross-section radius of gyration R_c_, obtained from rod-like Guinier analysis was ~13 Å, which is close to the disk thickness. Since the protein shape is more close to the sheet-like object (disk), R_c_ is used just as a reference here.

**Figure 3.**
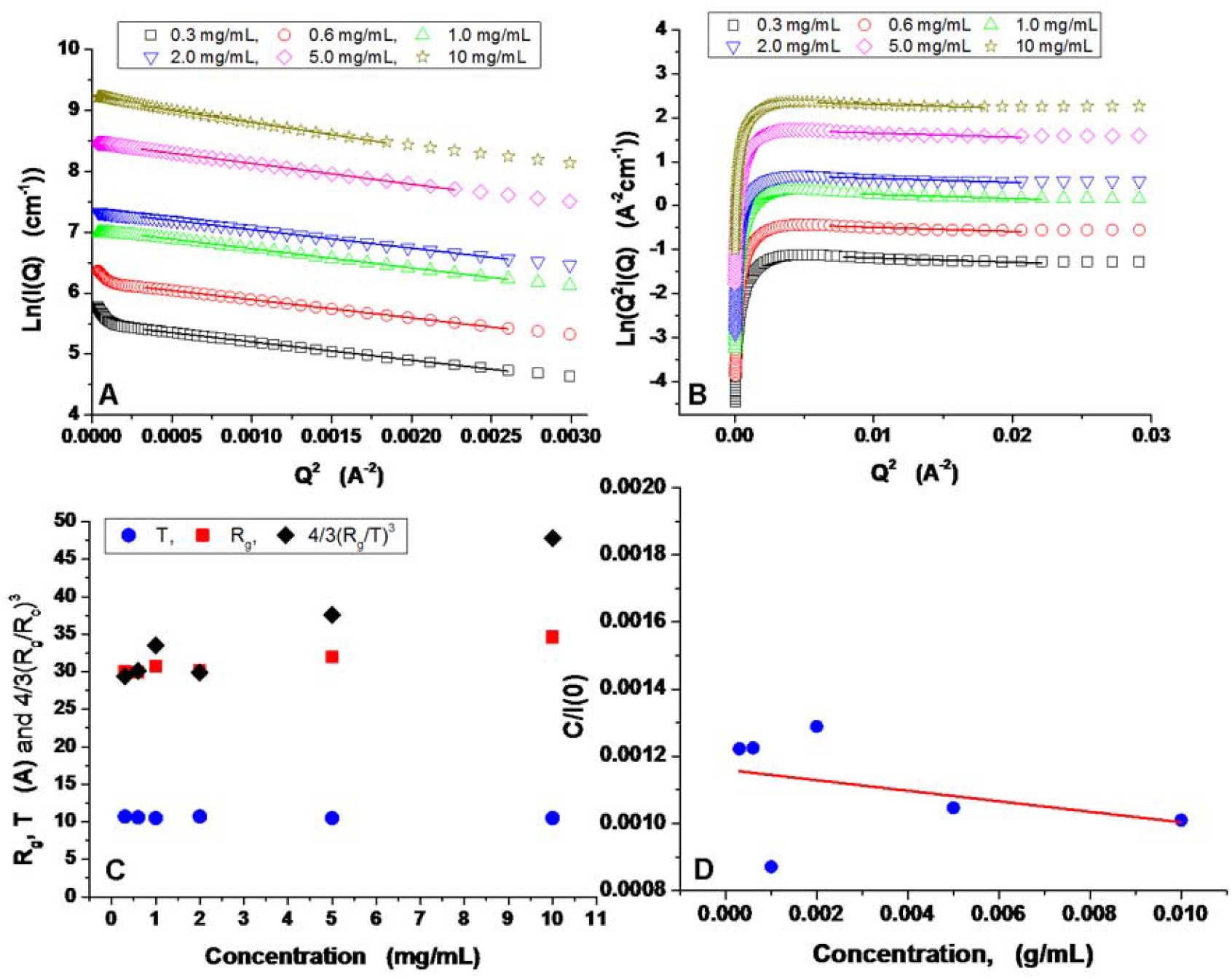
Guinier plots of (A) solid sphere and (B) disk for SAXS scattering intensity profiles of oat protein isolate in aqueous solution with various OPI concentration, (C) the concentration dependence of protein radius of gyration (R_g_), thickness (T) and 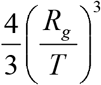, and (D) plot of C/I(0) as a function of protein concentration.

In addition to dimension (R_g_ and T), I(Q=0), the forward scattering was calculated during Guinier analysis, which is related to protein’s molecular weight and protein-protein interaction. Based on the forward scattering, we quantified the second virial coefficient (A_2_) of OPI in aqueous medium. The A_2_ calculation is based on the research of Tardieu et al.^37^, which assumed that proteins at low concentrations were under weak interactions. In detail, equation 1 was rewritten as I(Q)=cαP(Q)S(Q), where c is the protein concentration (mg/mL), and α, a pre-factor. The structure factor at q = 0 (S(0)) is related to the osmotic pressure, Δ, by 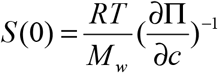, where *R* is the gas constant and *M_w_* is the molecular weight of protein. Π can be expressed through a virial expansion: 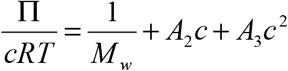… Higher order terms (i.e. *A_3_)* can be ignored, which generates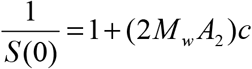. Based on the above derivation, we obtained 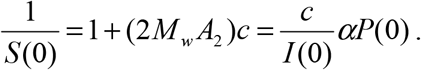 Hence, we plotted *c*/I(0) ~ *c*, and the intercept and slope were determined to be 0.00116 and −0.01566, respectively. From previous results of SDS-PAGE^43^, the OPI was composed of proteins or peptides with molecular weights of 66.2 kDa (2%), 57.0 kDa (3%), 45 kDa (3%), 36.0 kDa (38%), 22.0 kDa (43%), and 14.4 kDa (11%). Based on the composition, the number-averaged molecular weight of OPI was calculated to be 29.1 kDa (10^3^ g/mol). Then, the second virial coefficient (*A_2_*) of OPI in aqueous medium was determined to be −2.32×10^4^ mol*mL*g^−2^, which fell in the range that favored protein crystallization (−1×10^4^ to −8×10^4^ mol*mL*g^-2^)^44,45^. The osmotic second virial coefficient (*A_2_*) is a thermodynamic index that characterizes protein-protein interactions in dilute solutions by knowing the magnitude and sign of interaction^46^. Positive *A_2_* value suggests predominantly repulsive intermolecular interactions. Such attraction interaction between OPI particles may contributes to the aggregation of OPI at high concentrations^47^. The negative sign of second virial coefficient (*A_2_*) of OPI in aqueous medium indicated the dominant attractive interaction between OPI particles in aqueous medium. The *A_2_* of OPI in aqueous medium (-2.32×10^4^ mol*mL*g^-2^) also suggested a weak attractive interaction. Such weak attraction may be the driving force of the aggregation of OPI at high concentrations.

### 3.4. Effective Structure Factor

The effective structure factors (S_eff_) of oat protein isolate (OPI) in aqueous solution were calculated based on equation 5 and shown in **Figure 4A**, by referring to the scattering intensity profile at *C*_0_=0.3 mg/mL dilute solution. At low concentration (0.6~2.0 mg/mL), a peak at Q*~0.03 Å^-1^ showed up, suggesting the attractive interaction among proteins. When concentration reached high values (5 mg/mL and 10 mg/mL), the peak shifted to Q*~0.013 Å^-1^. The peak in S_eff_(Q) at Q* (finite Q) corresponds to the correlations between protein particles^48^. In other words, this Bragg peak from particle planes qualitatively indicates the mean nearest-neighbour particle distance *d* with *d*≈2π/Q*, also called correlation length^34^. **Figure 4B** displays the plot of correlation length as a function of protein concentration. At low concentration (0.6~2.0 mg/mL), the correlation length of oat protein was within 250 Å; when protein concentration increased to 5 mg/mL and 10 mg/mL, the correlation length immediately increased to more than 410 Å. The theoretical correlation length among proteins was calculated via 2(3*M*_w_/4π*N*_A_*C*)^1/3^ by taking 29.1 kDa as the molecular weight of oat protein isolate. From theoretical calculation, the increase of protein concentration should result in the decrease of correlation length. However, we observed that the concentration increase led to a rise of correlation length, which suggested the formation of large protein aggregates. It is reasonably speculated that once oat proteins form large aggregates, the inter-space between protein aggregates increased rather than decreased.

**Figure 4.**
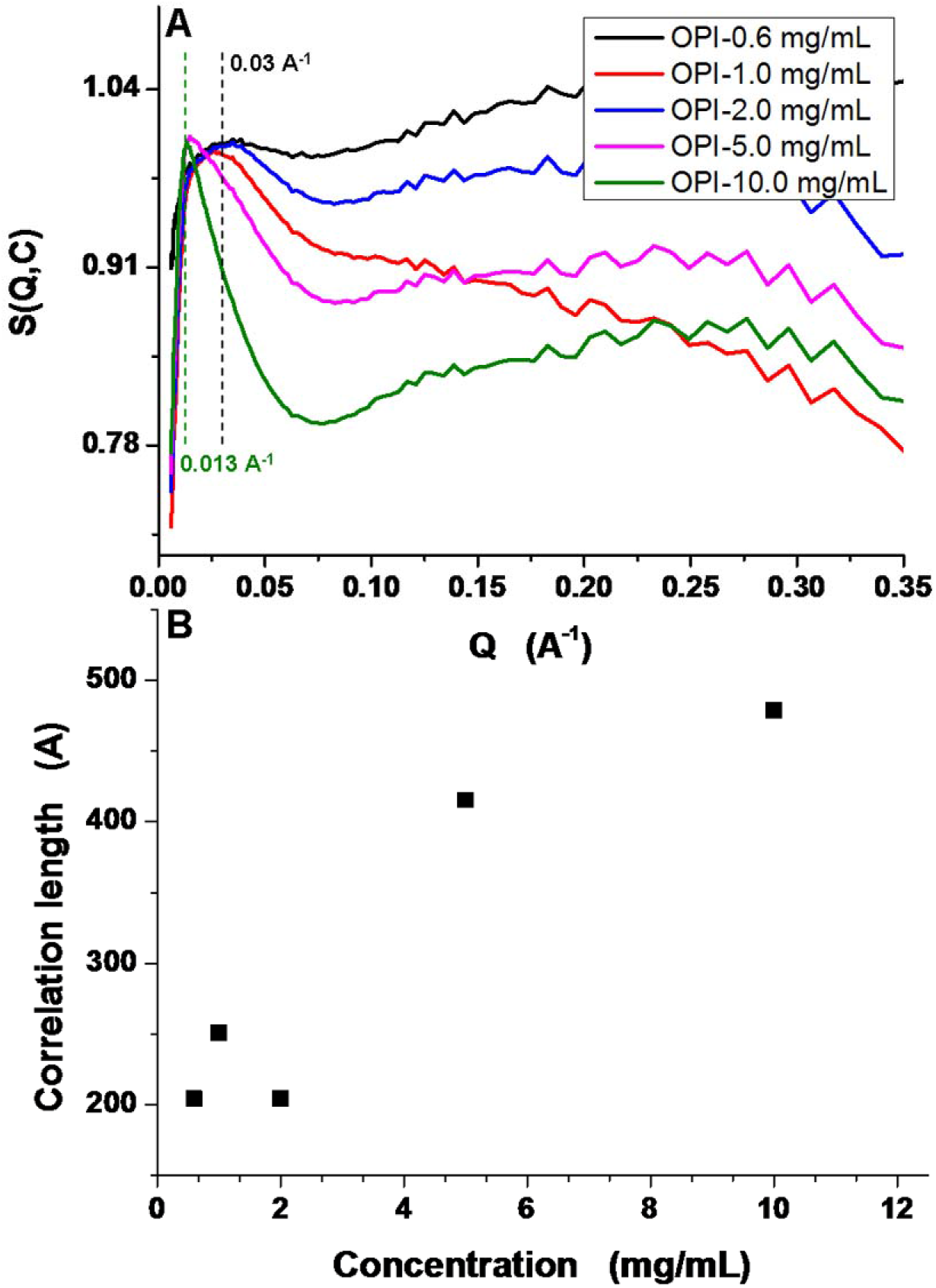
Effective structure factor analysis. (A) The effective structure factors S(Q,C) of OPI aqueous solutions at different concentrations from 0.6 mg/mL to 10.0 mg/mL, and (B) Plot of correlation length versus oat protein concentration.

### 3.5. Pair distribution function (PDF)

As a complimentary analysis, pair distribution function (PDF) (P(*r*)~*r*) is utilized to obtain the size distribution and *ab initio* contour restoration. The P(*r*) function indirectly provides information about the molecular shape and aggregation behavior, which allows more intuitive interpretation of the intensity profiles. The inverse transformation allows us to calculate the P(*r*) function based upon the equation: 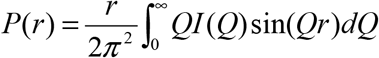. With P(*r*) function, we observed the point-point distance distribution within a protein molecule or aggregate of size *D_max_*. The algorithm GNOM was employed to extract the P(*r*) function curve from the intensity profile of I(Q)~Q^39^. **Figure 5** exhibits the P(*r*) curve of oat protein isolate (OPI) in aqueous solution from 0.3 mg/mL to 10 mg/mL. At low concentration from 0.3 mg/mL to 2 mg/mL, the P(*r*) curves for those three conditions almost overlapped with each other except for a little projection in the large Q region. The contours of P(*r*) curves confirmed the flatterned disk conformation of OPI particle with the feature of a distance at peak smaller than D_max_/2, by referring to the empirical scattering profiles and equivalent P(*r*) functions^38^. The fixed *D_max_* responded to the cross-section diameter of the individual protein disk. When protein concentration increased to 5 mg/mL and 10 mg/mL, the *D_max_* shifted from 80 Å to 100 Å and 110 Å, respectively. The contour of P(*r*) curve evolved into more skewed distributions with a peak position identical to those of low protein concentrations. The *D_max_* increase, unchanged peak position and more skewed contour suggested that high concentration induced a pseudo-ellipsoidal/rod-like aggregate in oat protein solution. Based upon the dimensions of P(*r*) functions, we speculated a concentration-dependent aggregation behavior for OPI in aqueous medium. It is worth-mentioning that different analytical methods result in different dimensional calculations. PDF is among those SAXS data analytical approaches.

**Figure 5.**
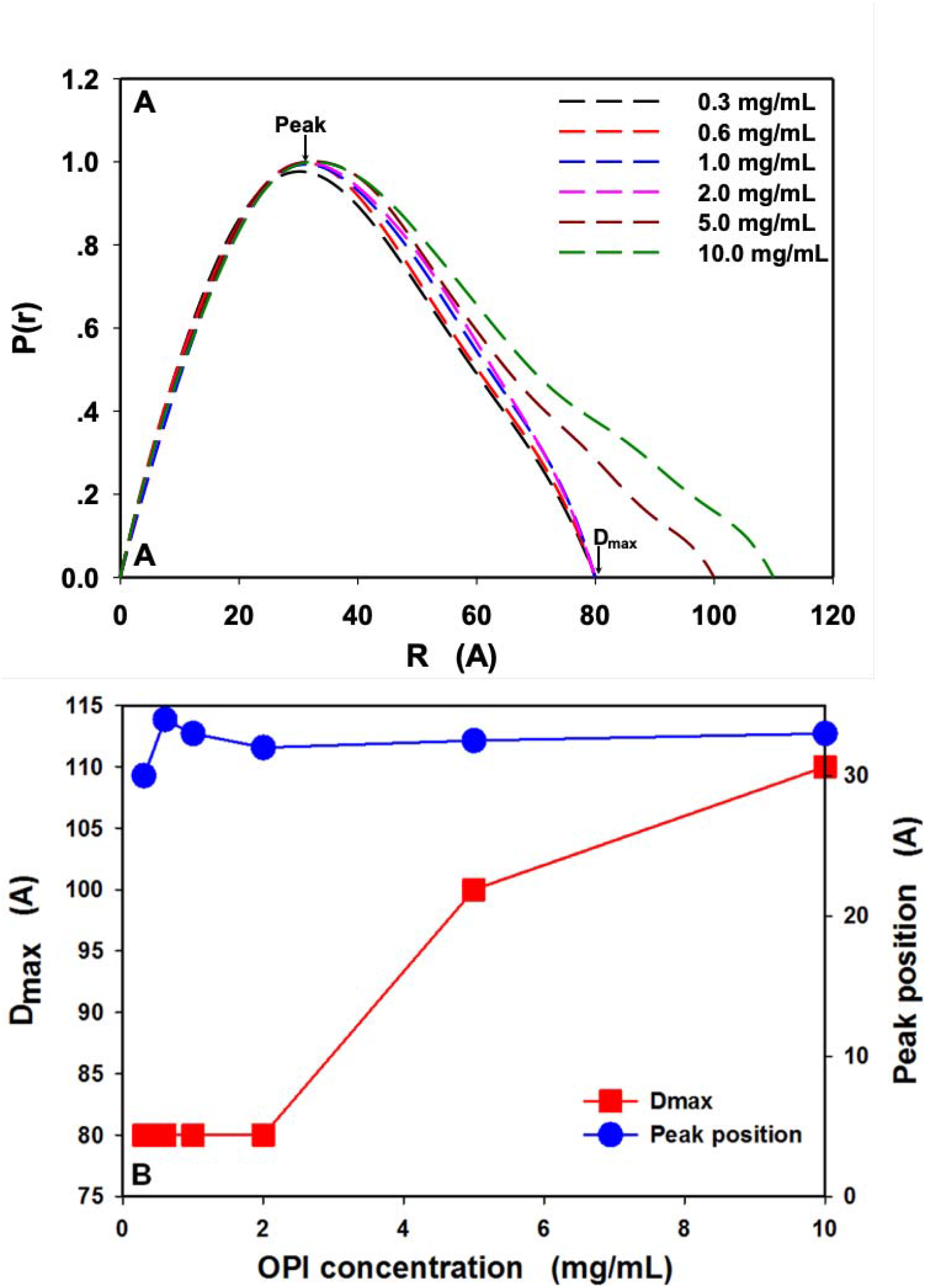
(A) Pair distribution functions of oat protein isolate in aqueous solution from 0.3 mg/mL to 10 mg/mL, (B) Plots of D_max_ and peak position as a function of OPI concentration.

### 3.6. *Ab initio* Restoration

To compare the envelopes of OPI in aqueous medium at different concentrations, *ab initio* models were restored based on the PDF results of OPI at 0.3 mg/mL, 0.6 mg/mL, by using algorithms GASBOR. This simulation approach is similar to DAMMIN or DAMMIF. The major difference lies in that protein structure is represented by an ensemble of dummy residues in continuous space. Besides, this approach runs much faster than others. 200 to 500 dummy residues were all used as simulation condition, which corresponds to molecular weight ranging from 20 kDa to 60 kDa. Since the OPI’s average molecular weight was close to 30 kDa, we chose the GASBOR models generated from the condition of 300 dummy residues. **Figure 6** shows the *Ab initio* restoration by using GASBOR dummy residue models of OPI at 0.3 mg/mL, 0.6 mg/mL, 1.0 mg/mL, and 2.0 mg/mL. For each condition, three different orientations are shown to give a comprehensive observation. From **Figure 6A, 6D**, we could clearly distinguish a thin disk contour for individual OPI particle at 0.3 mg/mL and 2.0 mg/mL. We also observed an ellipse or a mixture of ellipse and disk from **Figure 6B, 6C** for OPI particle at 0.6 mg/mL and 1.0 mg/mL. Now, it is safe for us to confirm the structural imaging of OPI from light scattering tool. The individual OPI particle forms thin disk or ellipse shape in physiological condtion.

**Figure 6.**
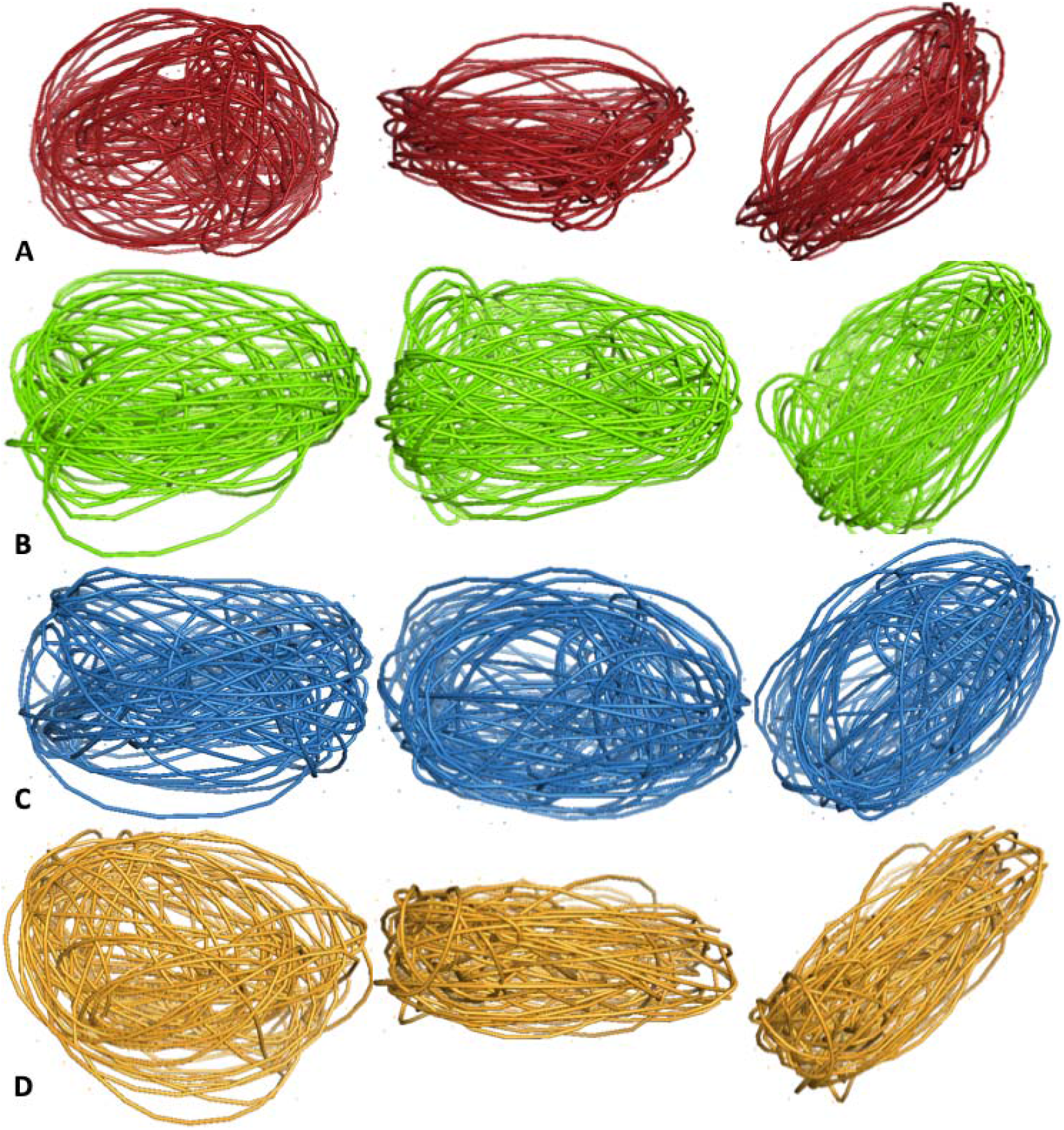
*Ab initio* restoration by using averaged GASBOR dummy residue model of OPI at (A) 0.3 mg/mL, (B) 0.6 mg/mL, (C) 1.0 mg/mL, and (D) 2.0 mg/mL. For each model, three different orientations are presented including original, rotate 90° along x axis, and rotate 45° along z axis (from left to right).

## 4. Conclusions

In summary, the structure analysis of oat protein isolate (OPI) in aqueous medium has been experimented by synchrotron small angle X-ray scattering and analyzed with multiple data approaches. The performance of structure experiments is within the pH window of human physiological condition. Through form factor analysis, we confirmed the disk conformation (41.4×41.4×10.2 Å^3^) of OPI in diluted aqueous mediums. With the increase of protein concentration (5 mg/mL and 10 mg/mL), OPI formed rod-like aggregates which was evidenced by the analyses of effective structure factor and pair distribution function (PDF). From PDF, we found that the peak position was independent on concentration increase, while as protein concentration increases to 5 mg/mL and 10 mg/mL, D_max_ increased from 80 Å to 100 Å and 110 Å, respectively. The PDF results suggested that the individual disk proteins aligned together to form rod aggregate upon protein concentration increase. Besides, the low-resolution *ab initio* protein models at the conditions from 0.3 mg/mL to 2.0 mg/mL were restored by using algorithms of GASBOR. We speculate that individual OPI most-likely forms thin disk shape at low concentration and it is a confirmation of our previous imaging observation.

## Acknowledgement

We thank Dr. Gang Liu for providing the oat protein isolate samples and BIO-CAT, ID18 beamline for performing SAXS experiment. This project was supported by the United States Department of Agriculture National Research Initiative (#2009-35603-05075).

